# *Leishmania* differentiation requires ubiquitin conjugation mediated by a UBC2-UEV1 E2 complex

**DOI:** 10.1101/2020.07.09.195024

**Authors:** Rebecca J. Burge, Andreas Damianou, Anthony J. Wilkinson, Boris Rodenko, Jeremy C. Mottram

## Abstract

Post-translational modifications such as ubiquitination are important for orchestrating the cellular transformations that occur as the *Leishmania* parasite differentiates between its main morphological forms, the promastigote and amastigote. 2 E1 ubiquitin-activating (E1), 13 E2 ubiquitin-conjugating (E2), 79 E3 ubiquitin ligase (E3) and 20 deubiquitinating cysteine peptidase (DUB) genes can be identified in the *Leishmania mexicana* genome but, currently, little is known about the role of E1, E2 and E3 enzymes in this parasite. Bar-seq analysis of 23 E1, E2 and E3 null mutants generated in promastigotes using CRISPR-Cas9 revealed numerous loss-of-fitness phenotypes in promastigote to amastigote differentiation and mammalian infection. The E2s UBC1/CDC34, UBC2 and UEV1 and the HECT E3 ligase HECT2 are required for the successful transformation from promastigote to amastigote and UBA1b, UBC9, UBC14, HECT7 and HECT11 are required for normal proliferation during mouse infection. Null mutants could not be generated for the E1 UBA1a or the E2s UBC3, UBC7, UBC12 and UBC13, suggesting these genes are essential in promastigotes. X-ray crystal structure analysis of UBC2 and UEV1, orthologues of human UBE2N and UBE2V1/UBE2V2 respectively, reveal a heterodimer with a highly conserved interface, highlighting the importance of stable UBC2-UEV1 interaction in the function of this complex across diverse eukaryotes. Furthermore, recombinant *L. mexicana* E1 UBA1a can load ubiquitin onto UBC2, allowing UBC2-UEV1 to form K63-linked di-ubiquitin chains *in vitro*. Notably, UBC2 can also cooperate *in vitro* with human E3s RNF8 and BIRC2 to form non-K63-linked polyubiquitin chains, showing that UBC2 can facilitate ubiquitination independent of UEV1, but association of UBC2 with UEV1 inhibits this ability. Our study demonstrates the dual essentiality of UBC2 and UEV1 in the differentiation and intracellular survival of *L. mexicana* and shows that the interaction between these two proteins is crucial for regulation of their ubiquitination activity and function.

**Author summary:** The post-translational modification of proteins is key for allowing *Leishmania* parasites to transition between the different life cycle stages that exist in its insect vector and mammalian host. In particular, components of the ubiquitin system are important for the transformation of *Leishmania* from its insect (promastigote) to mammalian (amastigote) stage and normal infection in mice. However, little is known about the role of the enzymes that generate ubiquitin modifications in *Leishmania*. Here we characterise 28 enzymes of the ubiquitination pathway and show that many are required for life cycle progression or mouse infection by this parasite. Two proteins, UBC2 and UEV1, were selected for further study based on their importance in the promastigote to amastigote transition. We demonstrate that UBC2 and UEV1 form a heterodimer capable of carrying out ubiquitination and that the structural basis for this activity is conserved between *Leishmania, Saccharomyces cerevisiae* and humans. We also show that the interaction of UBC2 with UEV1 alters the nature of the ubiquitination activity performed by UBC2. Overall, we demonstrate the important role that ubiquitination enzymes play in the life cycle and infection process of *Leishmania* and explore the biochemistry underlying UBC2 and UEV1 function.

## Introduction

Leishmaniasis is a neglected tropical disease caused by parasites of the genus *Leishmania*. This disease, which has a tropical and sub-tropical distribution, is transmitted by the bite of a sandfly and causes around 70,000 deaths annually (1). During their complex, digenetic life cycle, *Leishmania* differentiate between two main morphological forms: the motile, extracellular promastigote form in the sand fly vector and the non-motile, intracellular amastigote form in the mammalian host. Of the numerous promastigote forms that exist in the sandfly, procyclic promastigotes are the most actively dividing and metacyclic promastigotes are the transmissible form (2). In order for *Leishmania* cells to transition between the disparate environments of the sandfly mouthparts and mammalian phagolysosomes, substantial changes in gene expression, particularly at the post-transcriptional level, are required (3, 4). The post-translational modifications phosphorylation and ubiquitination, for example, are thought to contribute significantly to the differentiation process (5-8).

Ubiquitination regulates numerous cellular processes including proteasomal degradation, endocytic trafficking and DNA repair (9). Addition of the 8.5 kDa protein ubiquitin (Ub) to proteins is carried out by the sequential actions of E1 ubiquitin-activating (E1), E2 ubiquitin-conjugating (E2) and E3 ubiquitin ligase (E3) enzymes. Typically, an E1 activates ubiquitin in an ATP-dependent manner by adenylating its C-terminus, allowing a thioester bond to form between the E1 active site and ubiquitin. Subsequently, ubiquitin is transferred to the active site of an E2 via trans-thioesterification and then onto the substrate with the help of an E3 ligase. Most commonly, ubiquitination occurs on a lysine residue, although modification of cysteine, serine, threonine and the N-termini of proteins are also possible (10). E3 ligases can be grouped into two categories based on their mechanism of action. Cys-dependent E3s such as the HECT (Homologous to the E6-AP Carboxyl Terminus) and RBR (Ring-Between-Ring) E3s contain a cysteine residue that binds ubiquitin prior to transfer to the substrate. Conversely, Cys-independent E3s such as the RING (really interesting new gene) and U-box E3s facilitate the direct transfer of ubiquitin between E2 and substrate by providing a scaffold that orients the ubiquitin-charged E2 relative to the substrate (11). Ubiquitin-like modifiers (Ubls) such as SUMO and Nedd8 have a similar, but distinct, E1-E2-E3 conjugation pathway to that of ubiquitin (12). The removal of ubiquitin modifications is carried out by deubiquitinating enzymes (DUBs) (9).

A huge diversity of ubiquitin modifications exists, due in part to the ability of ubiquitin itself to be ubiquitinated on any of the epsilon amino groups of its 7 lysine side chains or on the alpha amino group of its N-terminus. This allows for the formation of linear or branched polyubiquitin chains. Acetylation, phosphorylation and the modification of ubiquitin with other Ubls (including SUMO and Nedd8) are also possible (13). The huge array of potential modifications permits a range of signalling outcomes. For example, K48-linked chains, the most common ubiquitin chain type, target proteins for proteasomal degradation. In contrast, K63-linked chains typically provide a non-degradative signal, for example in promoting the recruitment of proteins to sites of DNA damage (9).

To date, two ubiquitin-activating enzymes have been described in *Leishmania major* (14), and at least 20 cysteine peptidase DUBs exist in *Leishmania mexicana*, many of which are essential for the promastigote to amastigote transition (8). The importance of the ubiquitination system in these species is further exemplified by the finding that the *Leishmania* proteasome is essential for parasite survival, since many forms of ubiquitin modification target proteins for proteasomal degradation (15). The *L. major* Atg8 and Atg12 Ubls have been shown to play a role in parasite autophagy (16, 17) and, although they have yet to be properly described in *Leishmania*, SUMO- and Nedd8-conjugation systems exist in the closely-related kinetoplastid *Trypanosoma brucei* (18-23). Furthermore, a homologue of Urm1 is predicted to exist in *Leishmania* (24). However, little is currently known about the role that ubiquitin E1, E2 and E3 enzymes play in *Leishmania* biology. In this study we characterise the E1, E2 and E3 enzymes of *L. mexicana* by analysing the fitness of an E1, E2 and HECT/RBR E3 null mutant library throughout the *Leishmania* life cycle. The E2 ubiquitin-conjugating enzyme UBC2 and the ubiquitin E2 variant UEV1, both found to be essential for the promastigote to amastigote transition, are subsequently characterised in biochemical and structural detail.

## Results

### Ubiquitination gene families in *L. mexicana*

Initial analysis of the *L. mexicana* genome using Interpro and PFAM domain searches identified 4 ubiquitin-activating E1 (UBA), 15 ubiquitin E2-conjugating (UBC) and 81 E3 ligase genes (S1 Table). The putative E3s included 14 HECT, 1 RBR, 57 RING, 4 RING-CH-type and 5 U-box E3s. Upon more detailed analysis, however, LmxM.08.0220 and LmxM.02.0390 were found to be orthologues of *T. brucei* Uba2 and Ubc9, which have been identified as an E1 catalytic subunit and E2 enzyme for the Ubl SUMO respectively and were named UBA2 and UBC9 based on this orthology (21). Similarly, LmxM.01.0710 (UBA3) was found to share more similarity with HsUBA3 (68% query cover, 35.8% identity, E value: 8e^-64^), a Nedd8-activating enzyme catalytic subunit, than the ubiquitin E1 HsUBA1 (46% query cover, 31.7% identity, E value: 3e^-25^) and LmxM.24.1710 (UBC12) appears to be an orthologue of *T. brucei* Ubc12, an E2 Nedd8-conjugating enzyme (23). For two of the HECT domain-containing genes, *HECT13* and *HECT14*, only partial HECT domains of 69 and 60 amino acids were identified respectively, suggesting that they may be pseudogenes. Following the removal of Ubl E1 and E2s and possible pseudo-HECTs from the list of identified ubiquitination genes, 2 ubiquitin E1s, 13 ubiquitin E2s and 79 E3 ligase genes were proposed to be present in the *L. mexicana* genome. Of these, UBC1, a ubiquitin E2, is related to *T. brucei* CDC34 (99% query cover, 55.2% identity, E value: 2e^-105^), required for cytokinesis and infection progression of bloodstream form parasites in mice (25). UBC4, also a ubiquitin E2, is related to *T. brucei* PEX4 (100% query cover, 58.6% identity, E value: 5e^-72^), implicated in the ubiquitination of TbPEX5, a cytosolic receptor involved in peroxisome biogenesis (26).

Of the putative ubiquitination genes identified, the predicted molecular weights of encoded proteins range from between 115-127 kDa for E1s, 16-49 kDa for E2s, 147-733 kDa for HECT E3 ligases and 9-288 kDa for RING, RING-CH-type and U-box E3 ligases. Only one RBR-type E3 ligase, with a predicted molecular weight of 58 kDa, was identified. An alignment of UBA1a and UBA1b with human ubiquitin-activating E1 enzymes (S1A Fig) revealed conservation of the catalytic cysteine residue (C596 in UBA1a and C651 in UBA1b, equivalent to C632 in HsUBA1). Similarly, an alignment of *L. mexicana* and human E2-conjugating enzymes showed conservation of the conserved catalytic cysteine in all *L. mexicana* E2s except UEV1 (S1B Fig), suggesting the latter is a non-catalytic ubiquitin E2 variant (27). Furthermore, the HPN (His-Pro-Asn) motif, which is conserved in human E2-conjugating enzymes and which contains an asparagine residue that can be important for catalysis (28-30), was found to be partially or completely missing in UBC3, UBC6, UBC11, UBC14 and UEV1. A HECT domain alignment for *L. mexicana* HECT E3 ligases and 4 human HECT E3 ligases showed that HECTs 1-12 contained the conserved catalytic cysteine residue. Consistent with the identification of a partial HECT domain in HECT13 and HECT14, a putative catalytic cysteine was absent in these sequences (S1C Fig).

### A bar-seq screen reveals the importance of ubiquitination genes in the *L. mexicana* life cycle

In order to investigate the role of ubiquitination, SUMOylation and Neddylation genes in the life cycle of *L. mexicana*, the 4 E1, 15 E2, 12 HECT E3 and 1 RBR E3 genes were targeted for deletion in procyclic promastigotes using CRISPR-Cas9 (31). The resistance cassettes used for integration into the targeted gene locus contained unique, 12 nucleotide barcodes for the identification of individual null mutant lines in a pooled library. Null mutant lines were validated using PCRs to detect removal of the gene of interest and integration of the blasticidin resistance cassette (S2A Fig and S2B Fig) and, where a null mutant was not immediately generated, three rounds of transfection were performed to reduce the likelihood of not obtaining a null mutant line due to technical failure. A similar strategy was used for the generation of a Δ*hect12* mutant but will be described elsewhere. In this way, null mutants were generated for all genes except UBA1a, UBC3, UBC7, UBC12 and UBC13, suggesting that 1 out of 2 ubiquitin E1s and 4 out of 13 ubiquitin E2s may be essential in promastigotes. Although mutant clones were obtained for UBA1a, UBC3, UBC7 and UBC12, they still contained the gene of interest (S2C Fig). This lends further support to the proposition that UBA1a, UBC3, UBC7 and UBC12 are essential in promastigotes, since it suggests that gene duplication events have occurred to allow the parasite to retain the gene. Notably, a null mutant was successfully generated for the Nedd8 E1 UBA3 despite UBC12 (Nedd8 E2) appearing to be essential. This suggests that the essential role of UBC12 may be independent of UBA3. None of the SUMOylation, HECT E3 or RBR E3 genes were essential in promastigotes, as evidenced by the successful generation of null mutants for these genes.

Next, a bar-seq strategy, involving the parallel phenotyping of a null mutant pool using next-generation sequencing, was used to analyse the null mutants. Briefly, 58 promastigote-stage null mutants including 3 E1, 10 E2 (excluding Δ*ubc5*) and 13 E3 null mutants were pooled in equal proportions and in 6 replicate samples. Pooled promastigote (PRO) cultures were grown for 7 days and then induced to form axenic amastigotes (AXA) or used to purify metacyclic stage promastigotes (META). Purified metacyclic cells were used to infect macrophages (inMAC) in culture or mice using footpad injection (FP). At the time points indicated in Fig 1A, DNA was extracted to allow for amplification of the barcoded regions by PCR and quantitative analysis by next generation sequencing; the raw data are available in S4 Table in Damianou et al., 2020 (8). The heat maps in Fig 1B-D show the proportional representation of each null mutant line at each experimental time point, with a gradient of white to red representing proportions ranging between zero and 0.04 (or above) respectively. Proportional representation was calculated by dividing the number of reads for an individual barcode by the total number of reads for all expected barcodes and averaging over the 6 replicates. Loss-of-fitness phenotypes were inferred from decreases in proportional representation from one time point to the next. For Δ*uba2*, only 2.6 x 10^6^, instead of 4 x 10^6^ cells, were added to the pools due to the poor prior growth of this cell line. Furthermore, the associated barcode was not detected in all of the PRO 0 h time points, perhaps due to the lower representation of Δ*uba2* in the pool, prompting the decision to exclude this cell line from the analysis.

**Figure 1.**
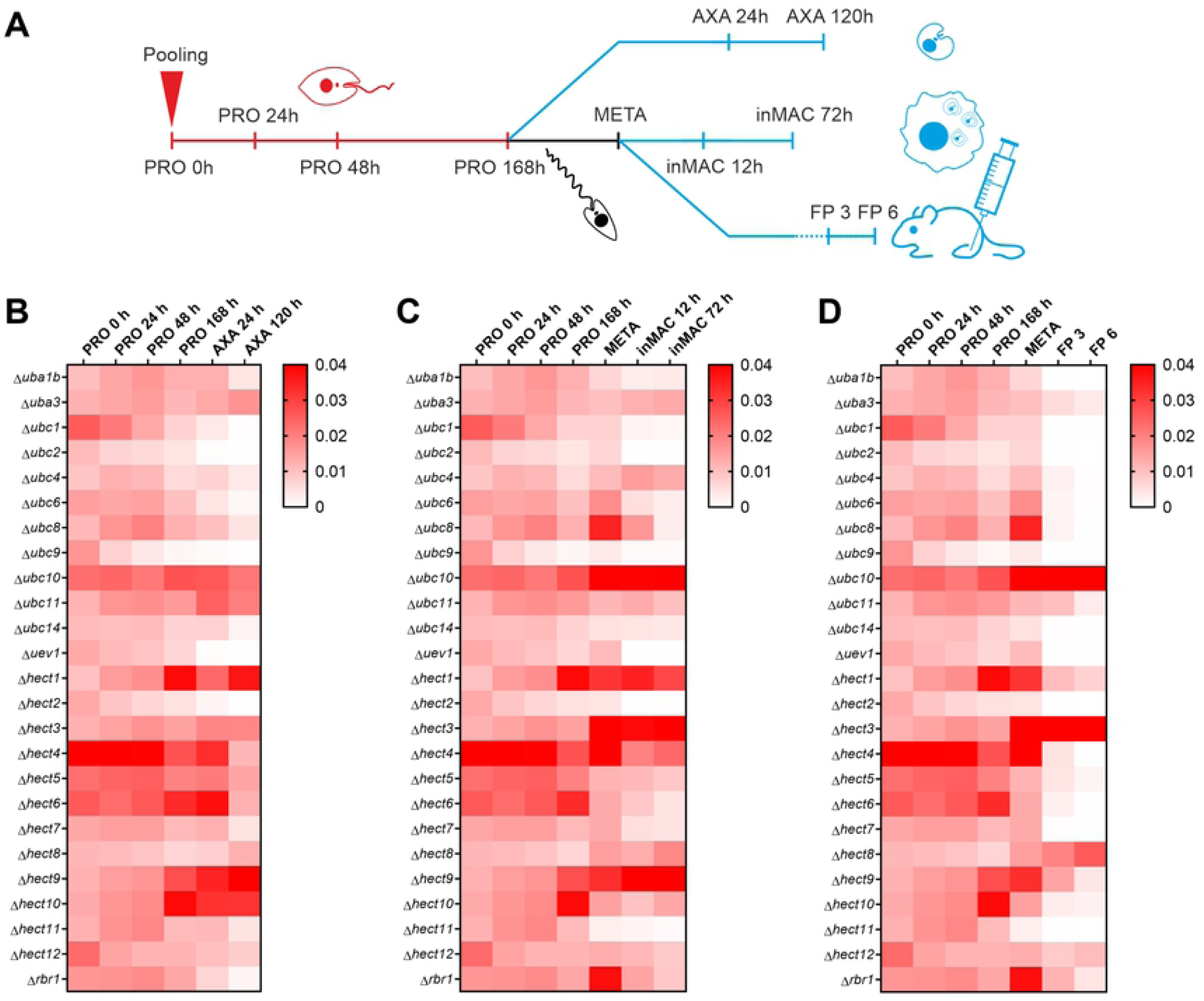
Life cycle phenotyping of ubiquitination gene null mutants. Fifty-eight null mutant lines were pooled (n = 6) as procyclic promastigotes and grown to stationary phase. Cells were then induced to differentiate into axenic amastigotes *in vitro* or metacyclic promastigotes purified and used to infect macrophages or mice. **A** Experimental workflow showing the time points at which DNA was extracted for barcode amplification and next generation sequencing. The heat maps for promastigote to **B** axenic amastigote, **C** macrophage infection or **D** mouse infection experiments show the average proportional representation of each null mutant at each experimental time point, calculated by dividing the number of reads for null mutant-specific barcodes by the total number of reads for expected barcodes. Samples included represent promastigote time-point zero (PRO 0 h), early-log phase (PRO 24 h), mid-log phase (PRO 48 h), late-log phase (PRO 72 h), stationary phase (PRO 168 h), early axenic amastigote differentiation (AXA 24 h), post-axenic amastigote differentiation (AXA 120 h), purified metacyclic promastigotes (META), early macrophage infection (inMAC 12 h), late macrophage infection (inMAC 72 h), 3 week footpad mouse infection (FP 3) and 6 week footpad mouse infection (FP 6).

Between the PRO 0 h and PRO 168 h samples, there was no complete loss of any null mutant line from the populations, although several significant loss-of-fitness phenotypes (decreases in proportional representation within the population) were observed between adjacent timepoints (<0.05, paired t-test, Holm-Šídák method, S4 Table, Damianou et al., 2020 (8)). For example, reduced fitness was seen at two or more promastigote time points for Δ*ubc9*, Δ*hect2* and Δ*hect12*. No loss-of-fitness phenotypes were observed between the PRO 168 h and META samples or between the PRO 168 h and AXA 24 h samples.

Within both the AXA 24 h-AXA 120 h and META-inMAC 12 h intervals, loss-of-fitness was observed for 10 null mutants: Δ*uba1b*, Δ*ubc2*, Δ*ubc6*, Δ*ubc8*, Δ*ubc9*, Δ*uev1*, Δ*hect2*, Δ*hect4*, Δ*hect7* and Δ*rbr1*, demonstrating good correlation between these two experiments. Of these, strong defects (>3-fold decrease in fitness between both intervals) were observed for Δ*ubc1/cdc34*, Δ*ubc2*, Δ*uev1* and Δ*hect2*. In addition, Δ*ubc4/pex4*, Δ*ubc11, Δubc14*, Δ*hect5*, Δ*hect6*, Δ*hect11* and Δ*hect12* showed loss-of-fitness between the AXA 24 h and AXA 120 h samples and Δ*hect10* showed a reduction in fitness between the META and inMAC 12 h samples. Further loss-of-fitness for Δ*ubc8* and Δ*rbr1* was observed between inMAC 12 h and inMAC 72 h. All null mutant lines that showed loss-of-fitness defects in the axenic amastigote and macrophage samples also had compromised fitness in the mouse, although many of the phenotypes observed were more severe. Specifically, Δ*uba1b*, Δ*ubc1/cdc34*, Δ*ubc2*, Δ*ubc9*, Δ*ubc14*, Δ*uev1*, Δ*hect2*, Δ*hect7* and Δ*hect11* had at least 20-fold reductions in fitness between the META and FP 3 samples. Additionally, Δ*uba3* and Δ*hect1* showed small decreases in fitness during this interval. Notably, Δ*ubc1/cdc34*, Δ*ubc2*, Δ*uev1* and Δ*hect2* showed the most dramatic and consistent loss-of-fitness phenotypes, including >5-fold, >25-fold and >200-fold decreases in fitness across the axenic amastigote, macrophage infection and mouse infection experiments respectively (calculated between the PRO 168 h/META sample and the experimental endpoints). A pBLAST search revealed that HECT2 is related to human UBE3C (28% query cover, 35.3% identity, 1e-64 E value), a HECT E3 ligase that ubiquitinates proteasome substrates to enhance proteasomal processivity (32). Overall, Δ*ubc2* and Δ*uev1* had the most severe phenotypes with reductions in fitness of 50-fold in the axenic amastigote, 300-fold in the macrophage and a complete loss-of-fitness (to zero) in the mouse. The strong defects observed for Δ*ubc1/cdc34*, Δ*ubc2*, Δ*uev1* and Δ*hect2* in pooled axenic amastigote differentiation were also observed individually using a cell viability assay (S2D Fig, S2 Table).

### UBC2 and UEV1 are orthologues of human UBE2N and UBE2V1/UBE2V2

The most probable orthologues of *L. mexicana* UBC2 (LmxM.04.0680) and UEV1 (LmxM.13.1580) were identified in the *Trypanosoma brucei, Saccharomyces cerevisiae* and human genomes using protein BLAST searches and the two groups of sequences aligned. Fig 2A shows that there is a high level of conservation between UBC2 and its predicted orthologues. Specifically, UBC2 shares 77%, 66% and 70% amino acid identity with the equivalent *T. brucei, S. cerevisiae* and human proteins respectively. Such a high level of shared identity suggests that UBC2 functions are likely to be well-conserved. In contrast, UEV1 is less well conserved with its orthologues, sharing 70%, 54% and 47% amino acid identity with the equivalent *T. brucei, S. cerevisiae* and human proteins respectively (Fig 2B). Consequently, there may be more functional divergence within this gene family. In humans, UBE2N and UBE2V1/V2 form a heterodimeric complex specific for K63-linked ubiquitination and have been linked to inflammatory signalling (33) and DNA damage response pathways (34).

**Figure 2.**
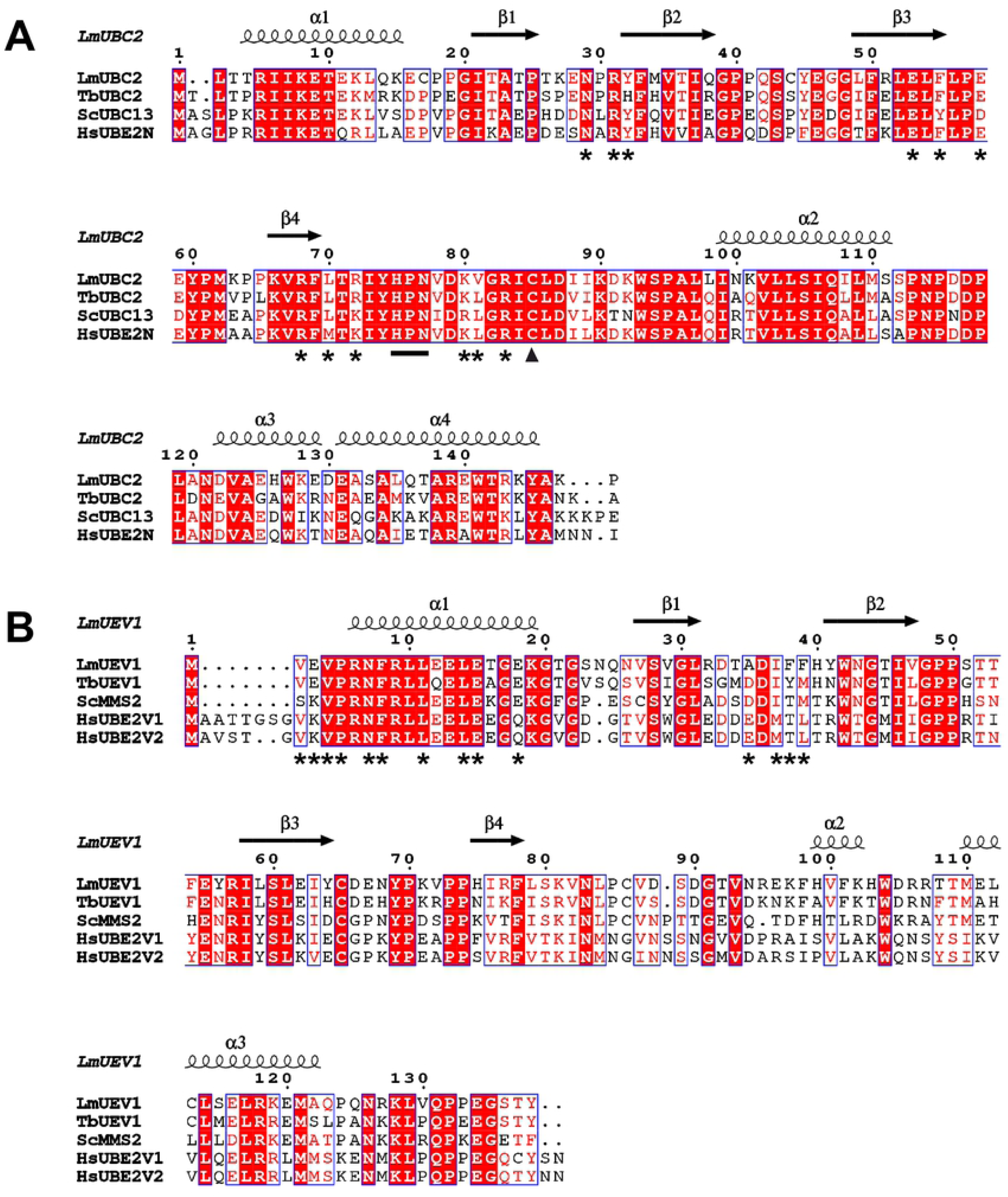
Alignments of LmUBC2 and LmUEV1 with selected orthologues. Sequence alignment and structural annotation were performed using T-Coffee and ESPript 3.0 respectively for **A** LmUBC2 and **B** LmUEV1. Red boxes indicate amino acid identity, red characters show similarity within the highlighted group and blue frames highlight similarity across groups. Positions of the UBC2 HPN motif and catalytic cysteine are shown by a black line and triangle respectively. Secondary structures derived from the crystallised *L. mexicana* UBC2-UEV1 heterodimer (Fig 4) are represented above the sequence alignment with helices represented by spirals and beta sheets by arrows. Asterisks denote important interface residues in the *L. mexicana* complex. *Lm, Leishmania mexicana; Tb, Trypanosoma brucei; Sc, Saccharomyces cerevisiae; Hs, Homo sapiens*.

*UBC2* and *UEV1* encode proteins of 148 and 138 amino acid residues respectively, corresponding to predicted molecular weights of 17 kDa and 16 kDa. Based on the alignment in Fig 2A, the putative catalytic cysteine of UBC2 is C85, equivalent to C87 in *S. cerevisiae* UBC13 and C87 in human UBE2N (35). UBC2 also contains an HPN motif 10 amino acids N-terminal of C85 that may be involved in catalysis (28-30). All of the proteins in the UEV1 alignment (Fig 2B) lack catalytic cysteine residues and HPN motifs, consistent with UEV1 being part of the noncatalytic UEV family of E2s (27). For the UEV1 alignment, isoform 3 (canonical sequence in UniProtKB) of UBE2V1 was used as this is the most similar isoform to UEV1. However, UBE2V1 is predicted to have numerous isoforms produced by alternative splicing. For example, a 30 residue N-terminal extension of isoform 2 (UEV-1A) relative to UBE2V2 has previously been shown to account for their differing functions (36). Since UEV1 lacks such an N-terminal extension, it could be predicted to be more similar in function to UBE2V2 than UBE2V1.

### *L. mexicana* UBC2 and UEV1 form a heterodimer

To permit biochemical characterisation of *L. mexicana* UBC2 and UEV1, His-Im9-tagged (37) UBC2 and UEV1 were expressed in *Escherichia coli* and purified following tag removal. In parallel, UBA1a, a putative ubiquitin-activating enzyme and the product of one of our proposed promastigote-essential genes, was similarly expressed and purified. Two putative ubiquitin-activating enzymes exist in *L. mexicana*, and, like *T. brucei* UBA1a and UBA1b, are more closely related to human UBA1 than UBA6 (14). Specifically, *L. mexicana* UBA1a and UBA1b share 36% and 33% amino acid identity with HsUBA1 respectively and 28% identity with each other. Based on its higher shared identity with HsUBA1 and potential essentiality, it was reasoned that UBA1a would be more likely to show a broad E2 specificity (comparable to that of HsUBA1) and therefore be capable of loading ubiquitin onto UBC2 (38, 39). All 3 proteins showed a good level of purity as assessed by InstantBlue staining (Fig 3A). For UBC2 and UEV1 no additional proteins were detected, whereas the UBA1a sample contained 2 extra proteins at around 74 and 100 kDa. These appeared below UBA1a, suggesting the presence of co-purified protein or UBA1a degradation products.

**Figure 3.**
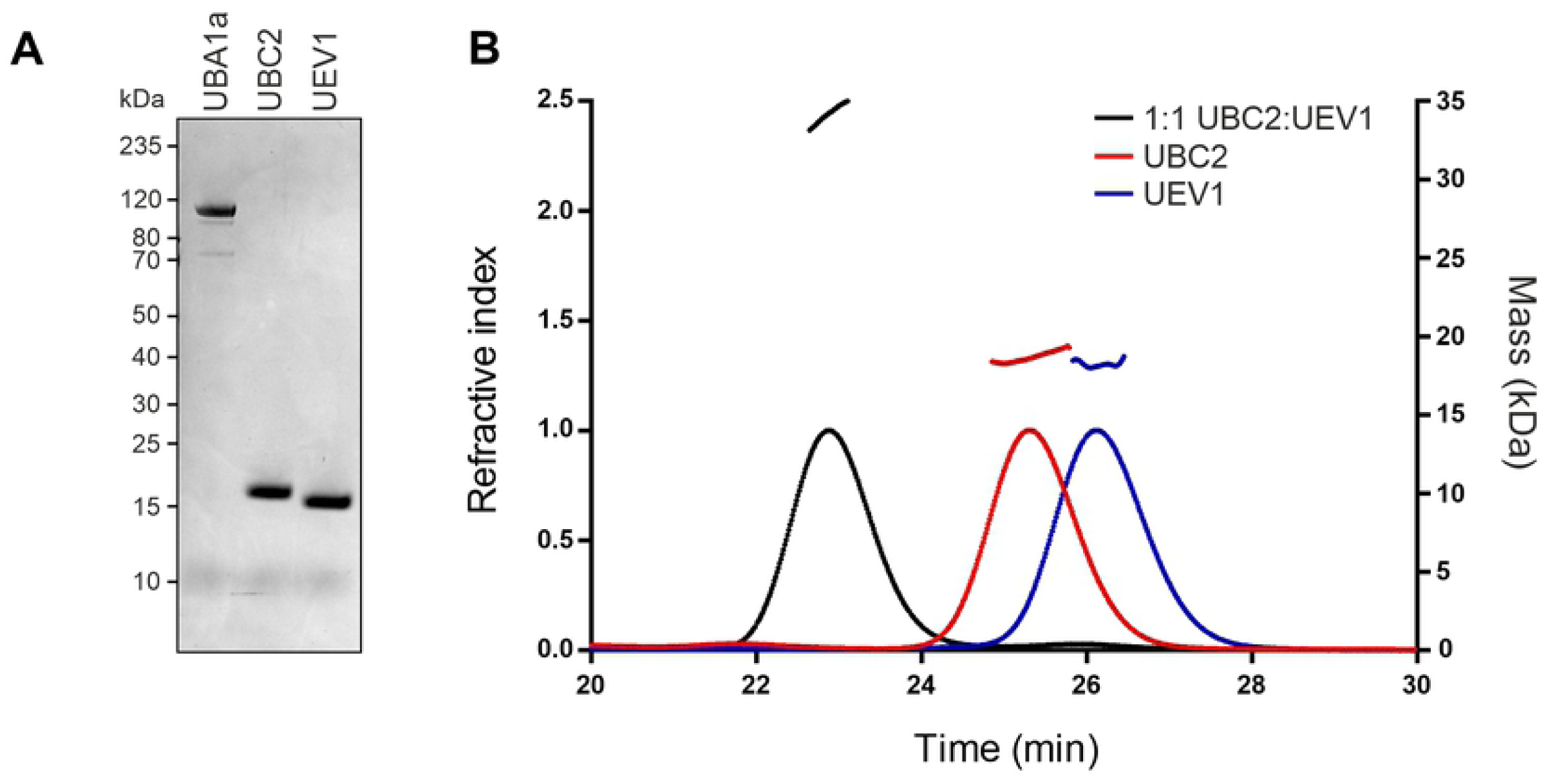
UBC2 and UEV1 form a stable heterodimer in vitro. **A** SDS-PAGE gel showing recombinant UBA1a, UBC2 and UEV1 stained with InstantBlue stain. Proteins were expressed in *E. coli* and purified by nickel affinity and size-exclusion chromatography. 1 µg of each protein was loaded onto the gel. **B** Elution profiles of UBC2, UEV1 and a 1:1 molar mix of UBC2-UEV1 presented as changes in refractive index over time. Curved lines show changes in refractive index for UBC2-UEV1 (black), UBC2 (red) and UEV1 (blue). The expected mass (as estimated from light scattering data) in kDa is indicated by a dashed line above the peak corresponding to each protein sample.

The orthologues of UBC2 and UEV1 in *T. brucei, S. cerevisiae* and humans have previously been shown to form a heterodimeric complex (40-43). To test whether this was also true in *L. mexicana*, a size-exclusion chromatography multi-angle laser light scattering (SEC-MALLS) approach was used. The chromatograms in Fig 3B show peaks in the refractive index representing the purified UBC2 and UEV1 samples at 25 and 26 min respectively. Measurement of both the refractive index and multi-angle laser light scattering allowed an estimation of the molecular weights of the recombinant proteins at 18.7 kDa and 18.2 kDa for UBC2 and UEV1 respectively. These values are close to the predicted molecular weights of 17 kDa for UBC2 and 16 kDa for UEV1. That both of these proteins eluted as a single peak is indicative of them existing in monomeric form while also reflecting the high quality of the protein preparations. When UBC2 and UEV1 were mixed in an equimolar ratio, an elution peak was seen at 23 min, indicating the presence of a higher molecular weight species (estimated at 34.2 kDa). Given that no peaks were seen representing UBC2 and UEV1 monomers, it can be assumed that all protein material in this sample existed in heterodimeric UBC2-UEV1 complexes. Therefore, *L. mexicana* UBC2 and UEV1 readily associate to form a heterodimeric UBC2-UEV1 complex.

### Structure of the UBC2-UEV1 heterodimer

To investigate the conservation and interactions of UBC2 and UEV1 at the structural level, we sought crystals of their complex. Crystals, which appeared after two days from polyethylene glycol-containing solutions, were sent for data collection at the Diamond Light Source, with the best crystal yielding a dataset extending to 1.7 Å spacing. The structure was solved by molecular replacement using the coordinates of the human UBE2N-UBE2V2 complex as the search model (PDB ID: 1J7D) (44). There are two UBC2-UEV1 heterodimers (PDB ID: 6ZM3) in the crystallographic asymmetric unit. These were found to have a similar structure and subunit organisation following superposition of chains equivalent to UBC2, UEV1 or the UBC2-UEV1 heterodimer by secondary structure matching (SSM) procedures, with RMSDs of 0.32 Å, 0.74 Å and 1.13 Å for 146, 132 and 276 equivalent atoms respectively. Since residues Gly20 to Asn24 are poorly defined in the electron density maps for one of the UEV1 chains, the alternative UBC2-UEV1 heterodimer was the focus of our analysis.

Like its *S. cerevisiae* and human orthologues, UBC2 has a canonical E2 structure comprising a 4-stranded antiparallel β-sheet flanked by four α-helices (Fig 4A). UEV1 exhibits a similar topology, but its polypeptide chain is shorter and the prominent pair of α-helices at the C-terminus of UBC2 (α3 and α4) are missing from UEV1. Relative to human UBE2V1 and UBE2V2, UEV1 has a shorter segment leading into the first α-helix (Fig 2B), a feature it shares with *S. cerevisiae* Mms2. Superposing UBC2 with human UBE2N, UEV1 with human UBE2V2 or the UBC2-UEV1 and UBE2N-UBE2V2 heterodimers (S3A Fig) gives RMSDs of 0.82, 1.15 and 1.13 Å for 147, 135 and 271 equivalent atoms respectively, demonstrating a high level of structural conservation between these orthologues. The conserved active site residues, His75, Pro76, Asn77 and Cys85, reside on an extended segment of the polypeptide that connects strand β4 and helix α2 of UBC2, with the thiol group of Cys85 projecting out of the catalytic cleft (Fig 4B). The high level of sequence conservation around these residues is reflected in the close proximity of their superposed catalytic clefts.

**Figure 4.**
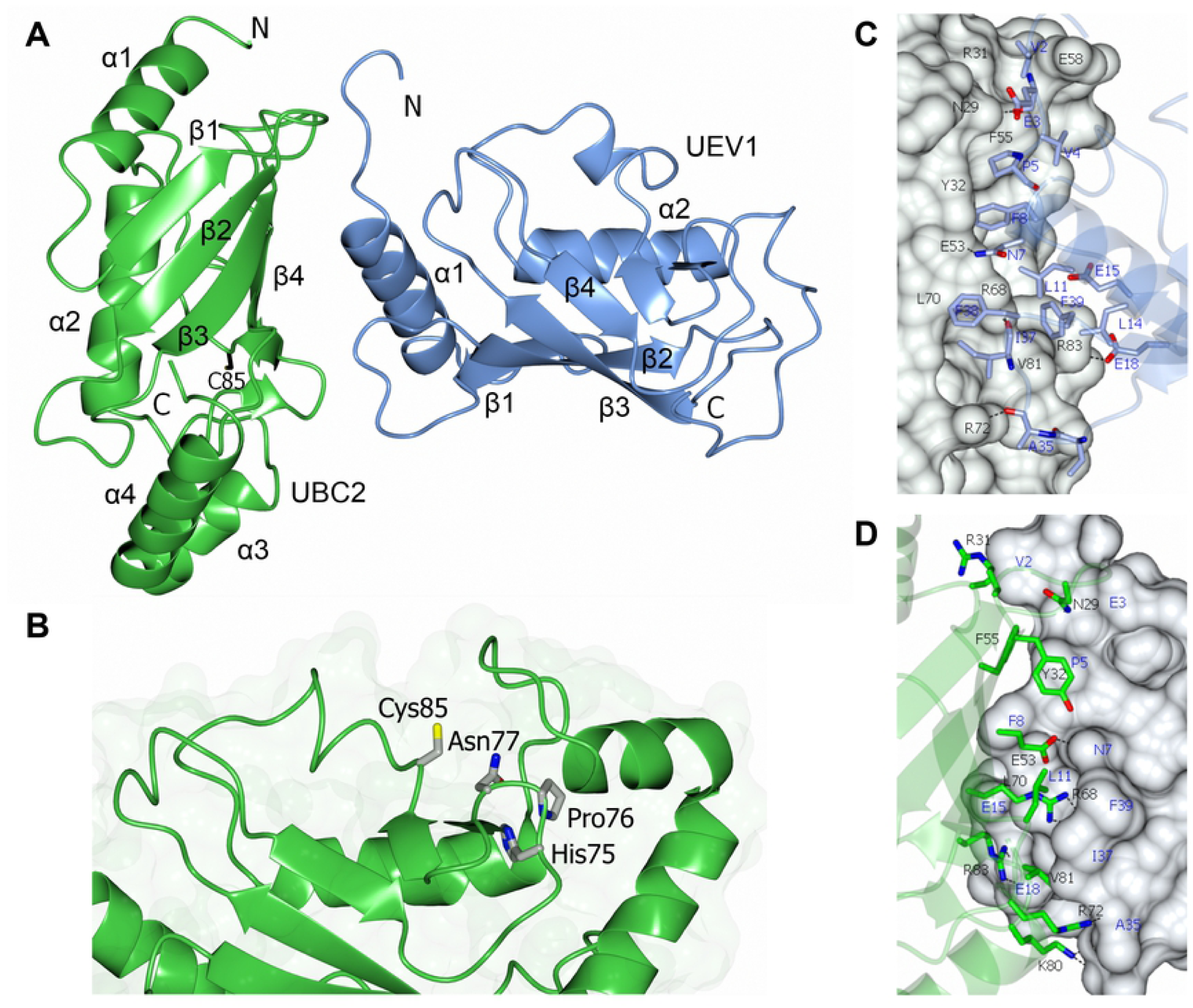
Structure of the UBC2-UEV1 heterodimer. **A** Ribbon diagram showing the crystal structure of UBC2 (green) and UEV1 (blue) in complex. The location of the N- and C-termini are highlighted along with the numbering of alpha helices (α) and beta strands (β). **B** Zoom-in of the conserved catalytic residues in UBC2. The HPN motif and proposed catalytic cysteine are shown as cylinders coloured by atom (red, oxygen; blue, nitrogen; yellow, sulfur). **C-D** Zoom-in of interface between UBC2 and UEV1 with UBC2 or UEV1 as a surface fill model respectively. Amino acid residues are shown as cylinders coloured by atom (red, oxygen; blue, nitrogen). Hydrogen bonds are denoted by dashed lines.

The interface between UBC2 and UEV1 (Fig 4C-D, S3B Fig) involves the β2, β3 and β4 strands and the loops following β1, β3 and β4 of UBC2 and the N-terminus, α1 helix and loop following β1 of UEV1. The buried surface area of this interface is 1,466 Å^2^. The core of the interface contains a number of hydrophobic residues (Tyr32, Phe55, Leu70 and Val81 of UBC2 and Pro5, Phe8, Leu11, Leu14 and Phe39 of UEV1) that contribute strongly to the interaction. Hydrophobic residues are highly conserved at these positions (Fig 2), highlighting the importance of the associated hydrophobic interactions for complex formation. These hydrophobic interactions are complemented by a number of polar intermolecular interactions, mostly notably between the side chains of Glu53 of UBC2 and Asn7 of UEV1 and Arg68 of UBC2 and the main chain carbonyl oxygen of Ile37 of UEV1. Both of these interactions are conserved in the human complex although Met is present in place of Ile. Notably, a salt bridge is formed between Arg83 of UBC2 and Glu18 of UEV1. Although these residues are conserved in *S. cerevisiae*, human UBE2V1/2 has Gln in place of Glu. Asp38 and Glu39 residues in human UBE2V2 provide hydrogen bonds for the interaction interface but are replaced by Thr and Ala (at positions 34 and 35) in UEV1 respectively.

### Ubiquitin transfer occurs between UBA1a and UBC2 *in vitro*

To assess whether UBA1a, UBC2 and UEV1 are an active E1, active E2 and inactive E2 variant respectively, recombinant proteins were tested in thioester intermediate assays. Fig 5A shows the results of incubating UBA1a, UBC2 and UEV1 in different combinations in the presence of ubiquitin and ATP. In these assays, human ubiquitin, which has 2 amino acid substitutions relative to *L. mexicana* ubiquitin (S4A Fig), was used. When UBA1a was present under non-reducing conditions, the appearance of a UBA1a∼Ub thioester intermediate (at around 125 kDa) was observed over time. Under reducing conditions, this intermediate was lost, confirming the presence of thioester-linked UBA1a∼Ub. When UBA1a and UBC2 were combined under non-reducing conditions, the appearance of an additional protein band at 26 kDa was observed, suggesting the transfer of ubiquitin between UBA1a and UBC2 to form thioester-linked UBC2∼Ub. This protein complex was lost under reducing conditions, confirming its identity as a UBC2∼Ub thioester intermediate. In contrast, when UBA1a and UEV1 were combined, no lower mobility species of UEV1 were observed, suggesting that UEV1 is unable to receive ubiquitin from UBA1a. When UBA1a, UBC2 and UEV1 were combined, no UBC2∼Ub thioester intermediate was observed. It was reasoned that UEV1 could prevent UBC2 from binding ubiquitin or, alternatively, that the presence of UEV1 facilitates the release of ubiquitin from UBC2 or its transfer onto substrate(s) in solution. Conversely, the alternative *L. mexicana* ubiquitin E1, UBA1b, was unable to transfer ubiquitin to UBC2, despite being capable of forming thioester-linked UBA1b∼Ub (S4B Fig). UBA1b was also unable to transfer ubiquitin onto UEV1 (S4B Fig), showing that ubiquitin cannot be loaded onto UEV1 by either of the two *L. mexicana* ubiquitin E1 enzymes. These results support the identities of UBA1a, UBC2 and UEV1 as an active E1, active E2 and inactive E2 variant respectively.

**Figure 5.**
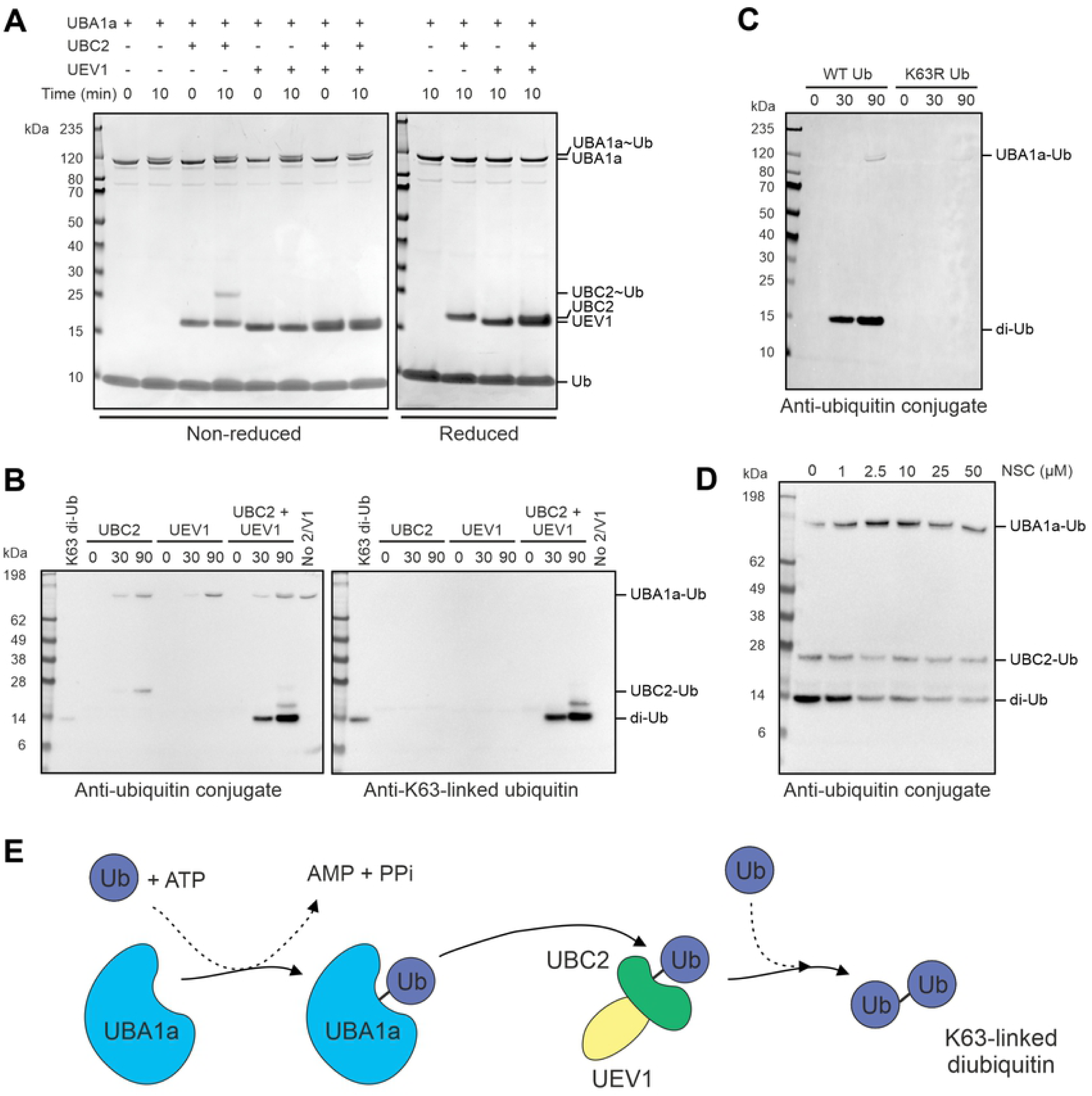
UBA1a and UBC2 cooperate in ubiquitin transfer in vitro. **A** Thioester assay demonstrating the ability of UBA1a and UBC2 to form thioester bonds with ubiquitin. UBA1a, ubiquitin and ATP were incubated with UBC2 and UEV1 as indicated in ubiquitination assay buffer for up to 10 min at room temperature. Samples were treated with either reducing or non-reducing sample buffer and visualised by SDS-PAGE with InstantBlue stain. **B** Di-ubiquitin formation assay. UBA1a, ubiquitin and ATP were incubated with UBC2 and UEV1 as indicated in ubiquitination assay buffer for up to 90 min at 37°C. Reactions were then treated with reducing sample buffer and visualised by immunoblotting with either ubiquitin conjugate or K63-linked ubiquitin antibodies. **C** Di-ubiquitin formation assay performed as for **B** but with wild-type ubiquitin substituted for K63R ubiquitin where shown. **D** UBC2 and UEV1 were pre-incubated with UBE2N inhibitor NSC697923 as indicated for 15 min prior to setting up a di-ubiquitin formation assay as in **B**. For **C** and **D**, samples were treated with reducing sample buffer prior to SDS-PAGE and immunoblotting with ubiquitin conjugate antibody. Data shown in **A-D** are representative. No 2/V1, no UBC2 or UEV1. **E** Schematic summarising the findings of **A-D**. UBA1a activates ubiquitin in an ATP-dependent manner, allowing ubiquitin to bind to the active site of UBA1a via a thioester linkage. Ubiquitin is then transferred to UBC2 (green) which, when present in complex with UEV1 (yellow), can generate free K63-linked ubiquitin chains.

### UBC2 and UEV1 conjugate ubiquitin *in vitro*

Previous *in vitro* studies have demonstrated the ability of *S. cerevisiae* UBC13 and MMS2 and their human counterparts to promote the formation of ubiquitin chains in the absence of E3 enzyme (28, 36, 41, 42, 45). To test whether the *L. mexicana* enzymes share this ability, different combinations of UBC2 and UEV1 were incubated with UBA1a, ubiquitin and ATP and di-ubiquitin formation monitored by immunoblotting for mono- and poly-ubiquitinated conjugates under reducing conditions. This approach was chosen in order to distinguish between UBC2, UEV1 and di-ubiquitin, which all have similar molecular weights. Di-ubiquitin formation was observed after 30 min in the presence of UBC2 and UEV1 and increased by 90 min (Fig 5B). In contrast, no di-ubiquitin formation was observed in the absence of either UBC2 or UEV1, suggesting that both proteins are required for di-ubiquitin formation. An additional protein that may represent tri-ubiquitin was observed above di-ubiquitin at the 90 minute time point, suggesting that higher order chains may be being assembled. Additionally, since the reducing conditions would disrupt any UBC2-Ub thioester intermediates, the protein at around 26 kDa in samples containing UBC2 may represent auto-ubiquitinated UBC2. Comparably, *in vitro* auto-ubiquitination of human UBE2N on K92, equivalent to K90 in UBC2, has been observed (42). Auto-ubiquitinated UBA1a also appears to be present at the top of the blot.

Since *S. cerevisiae* UBC13 and MMS2 and their human orthologues have been shown to specifically form K63-linked ubiquitin chains (41, 42), the reactions described above were additionally probed with a K63 linkage-specific antibody. The right-hand panel in Fig 5B shows that K63-linked ubiquitin conjugates are formed by UBC2-UEV1. Furthermore, UBC2 and UEV1 were unable to conjugate K63R mutant ubiquitin, demonstrating the essential requirement for K63 of ubiquitin in the formation of free ubiquitin chains by UBC2-UEV1 (Fig 5C). Pre-incubation of UBC2 with the covalent UBE2N inhibitor NSC697923 reduced di-ubiquitin formation in a concentration-dependent manner, showing that K63-linked ubiquitin chain formation is dependent upon UBC2 catalytic activity (Fig 5D). That NSC697923 was likely to inhibit UBC2 was rationalised based on the fact that all 4 of the residues that were mutated to make UBE2N resistant to NSC697923 are also found in UBC2 (46). Complete inhibition of di-ubiquitin formation was not achieved, perhaps due to an insufficiently long incubation time for UBC2 with NSC697923. These experiments demonstrate that UBC2 and UEV1 can form free K63-linked ubiquitin chains *in vitro* (Fig 5E).

The crystal structure of *S. cerevisiae* Mms2-Ubc13 covalently linked to a donor ubiquitin molecule revealed the structural basis of K63 linkage specificity in ubiquitin chain formation. In this structure, Mms2 directs the K63 residue of a putative acceptor ubiquitin into the Ubc13 active site, where it can attack Gly76 of the donor ubiquitin bound to the Ubc13 active site cysteine to form an isopeptide bond (47). An overlay of this structure (PDB ID: 2GMI) with chains A and B of our UBC2-UEV1 structure allowed the positions of the donor and acceptor ubiquitins to be revealed in the context of the *L. mexicana* complex (S3C Fig). This produces a plausible model of the quaternary complex without significant steric clashes and suggests that the same strategy is used to confer Lys63-linkage specificity in *L. mexicana* and *S. cerevisiae*. In support of this, two Mms2 residues shown to be required for acceptor ubiquitin binding in the *S. cerevisiae* Mms2-Ubc13-Ub structure, Ser27 and Thr44, are conserved in *L. mexicana* UEV1 (S28 and Thr45).

### UBC2 can cooperate with human E3s to allow polyubiquitination *in vitro*

As the cognate E3s for UBC2 and UEV1 have not yet been identified, human E3s known to catalyse ubiquitination in coordination with UBE2N were tested for their ability to similarly cooperate with UBC2 and/or UEV1. The reasons for doing this were threefold. Firstly, to investigate whether UBC2 could carry out the typical E2 role of facilitating ubiquitin transfer to substrates via E3 enzymes. Secondly, in the hope of making inferences about *L. mexicana* E3s that could be part of the UBC2 ubiquitination cascade and, lastly, to explore the conservation of E2-E3 interactions between *L. mexicana* and humans. For this purpose, two RING E3 ligases, BIRC2 and RNF8, were selected for testing on the basis that they are known to interact with human UBE2N (48-50). In humans, BIRC2 has wide-ranging roles including in regulating apoptosis and cell proliferation (48, 51) and RNF8 has well-characterised roles in DNA damage signalling (52-54). HUWE1, a HECT E3 ligase that is not known or predicted to interact with UBE2N, was also chosen for comparison (55). All recombinant E3s used were GST-tagged and, with the exception of HUWE1 which was N-terminally truncated, full-length.

Fig 6 shows the ubiquitination profiles observed for BIRC2, RNF8 or HUWE1 incubated with different combinations of UBC2 and UEV1 in reactions containing UBA1a, ubiquitin and ATP. When BIRC2 or RNF8 were incubated with UBC2 in the absence of UEV1, a prominent pattern of polyubiquitination was seen (upper panel). Based on the K63-linked ubiquitin and GST blots (middle and lower panels respectively), the polyubiquitination observed was not K63-linked and occurred both as a result of E3 auto-ubiquitination and of free chain formation and/or ubiquitination of other proteins (such as UBA1a, UBC2 or UEV1) in solution. When BIRC2 or RNF8 were incubated with UEV1 in the absence of UBC2, polyubiquitination did not occur. However, a single protein was present in the ubiquitin conjugate blot at around 130 kDa, likely representing ubiquitinated UBA1a (as observed in Fig 5B-D). Similarly, polyubiquitination was not observed when BIRC2 or RNF8 were incubated with both UBC2 and UEV1. In these samples, however, di-ubiquitin formation was notably increased, suggesting that UEV1 is able to regulate UBC2 such that it switches its activity between facilitating polyubiquitination by E3s and forming K63-linked ubiquitin chains in complex with UEV1 (Fig 6B). In contrast, polyubiquitination was observed when either UBC2, UEV1 or UBC2 and UEV1 were combined with HUWE1, suggesting that physical interaction between E2s and HUWE1 may be required for HUWE1 activity. Alternatively, the truncated nature of the recombinant HUWE1 protein, which lacks its UBA and WWE protein-protein interaction domains, may have encouraged non-specific interaction with and ubiquitin transfer from UBA1a. No ubiquitin conjugates were observed in the absence of both UBC2 and UEV1.

**Figure 6.**
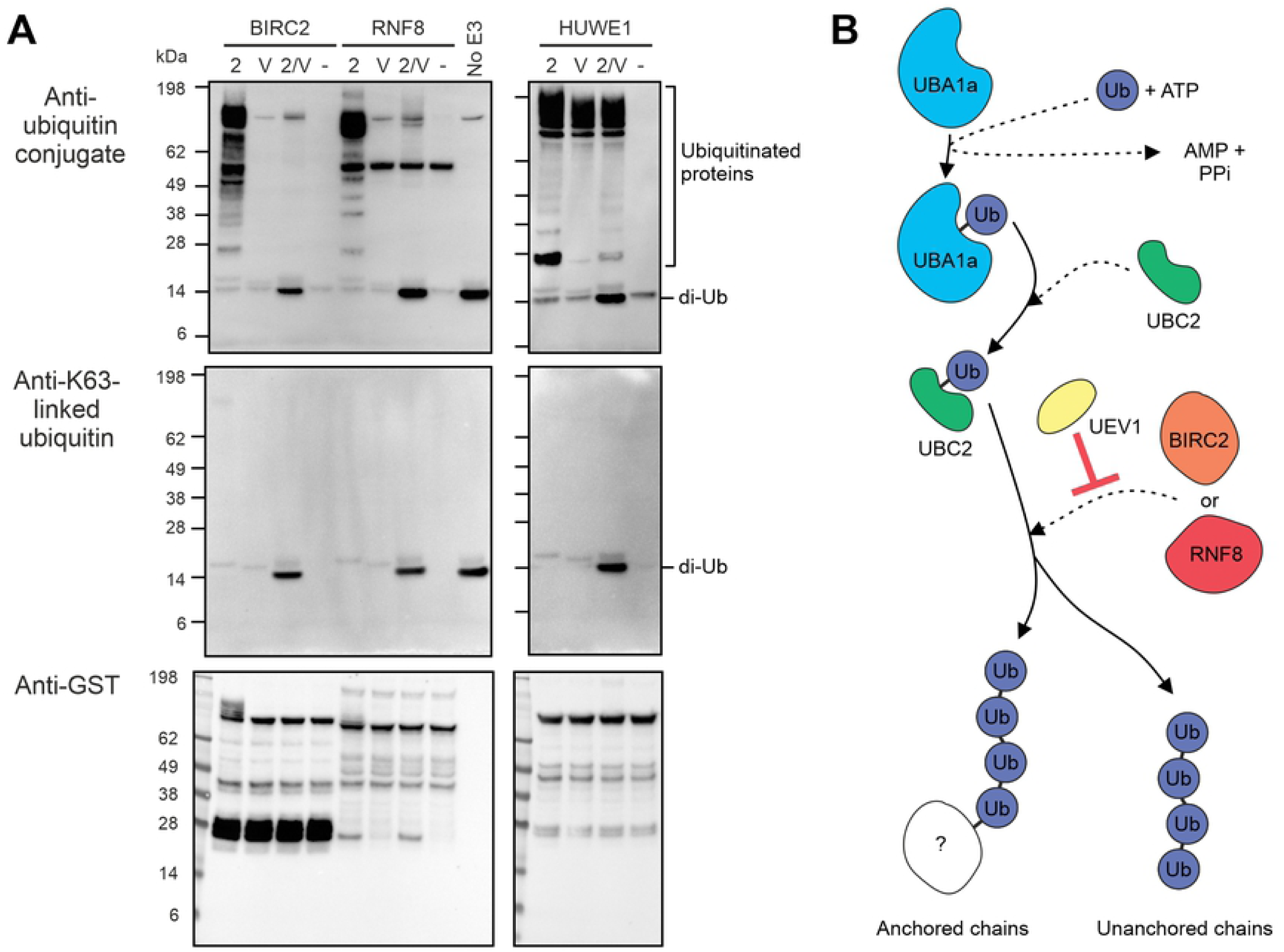
Cooperation of UBC2 with human E3s in in vitro polyubiquitination. **A** UBA1a, ubiquitin and ATP were incubated with UBC2, UEV1 and human E3s (BIRC2, RNF8 and HUWE1) as indicated in ubiquitination assay buffer for 1 h at 30°C. Reactions were visualised by immunoblotting with either ubiquitin conjugate, K63-linked ubiquitin or GST antibodies as shown. 2, UBC2; V1, UEV1; -, no UBC2 or UEV1; no E3, no BIRC2, RNF8 or HUWE1. **B** Schematic summarising the findings of **A**. UBA1a activates ubiquitin in an ATP-dependent manner, allowing ubiquitin to bind to the active site of UBA1a via a thioester linkage. Ubiquitin is then transferred to UBC2 which can, by interacting with the human E3s BIRC2 and RNF8, form non-K63-linked polyubiquitin chains. UEV1 inhibits this association, presumably via its interaction with UBC2.

### High functional conservation of human and *Leishmania* enzymes

In order to further interrogate the functional conservation between *Leishmania* and human enzymes, the ability of UBC2 to extend ubiquitin chains on the human U-box E3 ligase CHIP was investigated. *In vitro* monoubiquitination of CHIP by UBE2W can be followed by the extension of ubiquitin chains by UBE2N-UBE2V1 and provides an example of a pair of cooperating E2s with distinct chain initiation and elongation functions (56-58). To simplify the experimental setup and interpretation of results, it was decided to select a single E1 enzyme to facilitate ubiquitin transfer to both human UBE2W and *L. mexicana* UBC2. In a thioester assay, *L. mexicana* UBA1a was shown to be equally competent at transferring ubiquitin to UBE2W as human UBA1 (Fig 7A) and was therefore selected for use in subsequent experiments. In addition to the reducible UBE2W-Ub thioester bands observed, an additional, non-reducible band was observed following incubation of E1 and E2 enzymes. This is likely to be N-terminally ubiquitinated UBE2W (59, 60).

**Figure 7.**
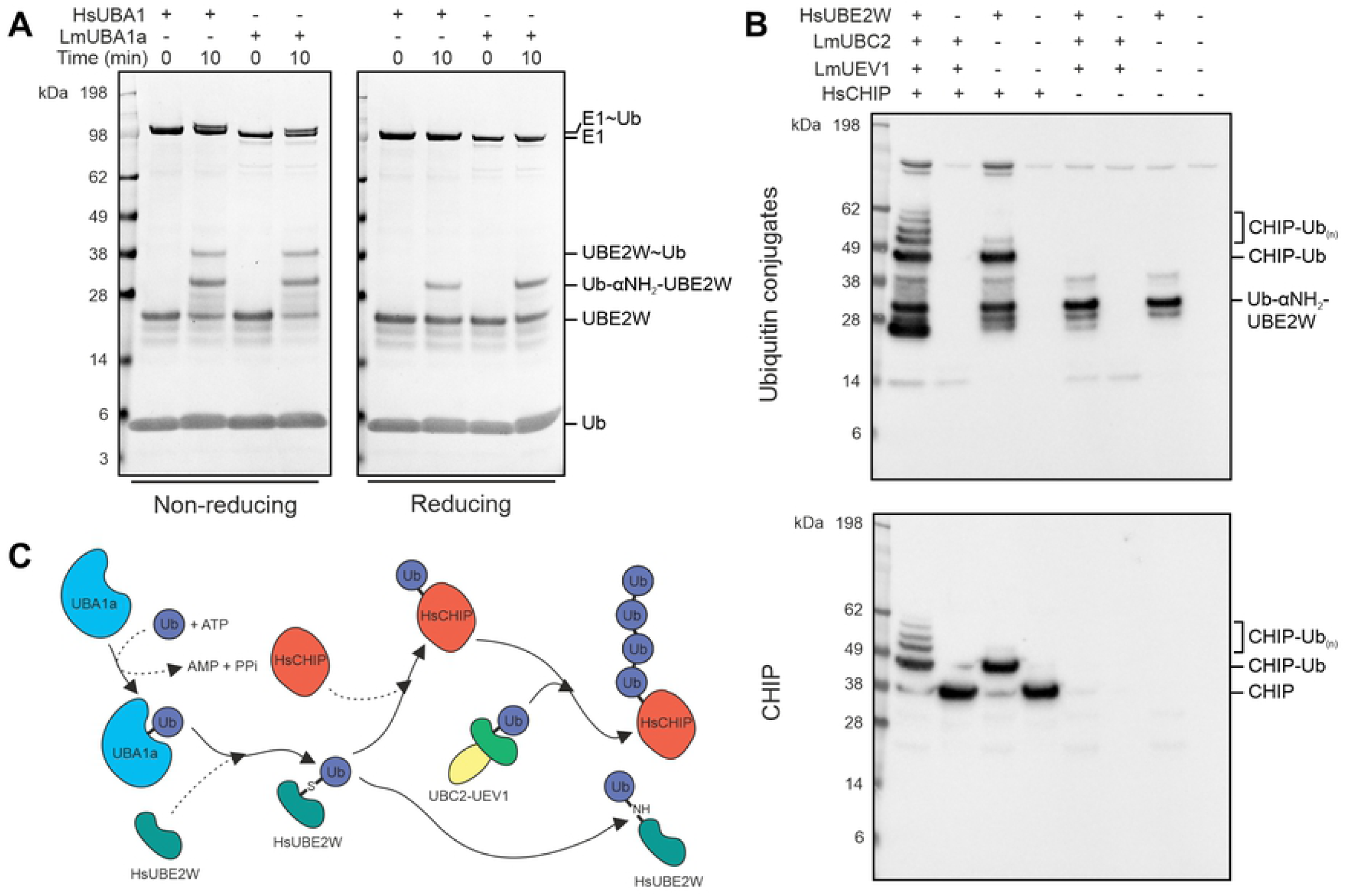
Ubiquitin chain extension activity of UBC2-UEV1. **A** Thioester assay showing ubiquitin transfer to UBE2W from both human UBA1 and *L. mexicana* UBA1a. UBE2W, ubiquitin and ATP were incubated with human UBA1 or *L. mexicana* UBA1a as indicated in ubiquitination assay buffer for up to 10 min at room temperature. Samples were treated with either reducing or non-reducing sample buffer and visualised by SDS-PAGE with InstantBlue stain. **B** UBA1a, ubiquitin and ATP were incubated with UBC2, UEV1, human UBE2W and human CHIP as indicated in ubiquitination assay buffer for 1 h at 30°C. Reactions were visualised by immunoblotting with either ubiquitin conjugate or CHIP antibodies as shown. **C** Schematic summarising **A-B**. UBA1a activates ubiquitin in an ATP-dependent manner, allowing ubiquitin to bind to the active site of UBA1a via a thioester linkage. Ubiquitin is then transferred to HsUBE2W which monoubiquitinates HsCHIP. Alternatively, HsUBE2W can ubiquitinate its own N-terminus. Once primed with monoubiquitin, UBC2-UEV1 can extend ubiquitin chains on HsCHIP. Lm, *Leishmania mexicana*; Hs, *Homo sapiens*. Where species is not indicated, proteins are from *L. mexicana*.

When *L. mexicana* UBC2 and UEV1 were incubated with human UBE2W and CHIP in the presence of E1, ubiquitin and ATP, polyubiquitinated CHIP was observed (Fig 7B). When UBE2W was absent from this reaction, no CHIP ubiquitination or free chain formation was seen. Alternatively, when UBC2 and UEV1 were absent, only monoubiquitinated CHIP was observed, suggesting that UBE2W is priming CHIP with a single ubiquitin modification that can then be extended by UBC2 and UEV1 (Fig 7C). In this respect, UBC2 and UEV1 behave in a similar manner to human UBE2N-UBE2V1 (56). The absence of free chain formation in the presence of UBC2, UEV1 and CHIP, however, is in contrast to what was reported for human UBE2N-UBE2V1 and CHIP (56, 57). In reactions where UBE2W was present but CHIP was absent, a strong band at around 34 kDa was observed, likely corresponding to N-terminally ubiquitinated UBE2W as in Fig 7A (59, 60).

## Discussion

Trypanosomatids are amongst the most ancient of eukaryotes and possess some highly divergent biochemistry, for example compartmentalisation of the glycolytic pathway or mRNA trans-splicing (61, 62). Despite this, our bioinformatic analysis of the *L. mexicana* genome revealed numerous ubiquitin conjugation system components: 2 ubiquitin E1, 13 ubiquitin E2 and 79 E3 ligase genes, including 12 HECT E3s, 1 RBR E3, 5 U-box RING E3s and 4 RING-CH-type E3s. These numbers are similar to those of another single-celled eukaryote, *S. cerevisiae*, which has 1 E1, 11 E2s and 60-100 E3s, of which 5 are HECT E3s, 2 are RBR E3s, 2 are U-box E3s and at least one is a RING CH-type E3 (63, 64). In contrast, humans have 2 E1s, 40 E2s and over 600 E3s, of which 28 are HECT E3s, around 15 are RBR E3s, 9 are U-box E3s and 11 are RING CH-type E3s (64-68). These numbers reveal that *Leishmania* may have a higher ratio of HECT:total E3s than *S. cerevisiae* or humans due to an expansion of this gene family in the *Leishmania* lineage. We also identified a putative SUMO E1 catalytic subunit (UBA2), a Nedd8 E1 catalytic subunit (UBA3), a SUMO E2 (UBC9) and a Nedd8 E2 (UBC12), although E1, E2 and E3 genes for Ubls were not extensively searched for. Despite this, previous characterisation of the *T. brucei* orthologues of UBA2 and UBC9 by *in vitro* SUMOylation assays and UBA3 and UBC12 by affinity purification of Nedd8 lend further weight to their proposed functions (21, 23). Of the putative ubiquitin E2s identified, 5 were missing the conserved asparagine residue thought to be important for E2 catalysis (28-30). UEV1, which also lacks the catalytic cysteine residue, is part of the non-catalytic ubiquitin E2 variant protein family. For UBC3, UBC6, UBC11 and UBC14, in contrast, the missing asparagine residue may indicate non-canonical methods of E2 catalysis.

Previous work has underscored the importance of the ubiquitination system in *Leishmania* by demonstrating both the essentiality of the parasite proteasome and a key role for DUBs in the life cycle of this parasite (8, 15). Our generation of a select ubiquitination gene null mutant library revealed that 1 out of 2 ubiquitin E1s and 4 out of 13 ubiquitin E2s could not be deleted and therefore may be essential in promastigotes. A smaller proportion (4 out of 20) cysteine peptidase DUBs were shown to be essential in promastigotes, perhaps due to a greater degree of redundancy in DUB function (8). UBC12, a putative Nedd8 E2, also appears to be essential in promastigotes, suggesting it functions independently of the Nedd8 E1 UBA3, which is non-essential in promastigotes. This could be due to the presence of an additional Nedd8 E1 or a Neddylation-independent function of UBC12. In contrast, none of the HECT or RBR E3 ligases were essential in promastigotes, likely attributable to the considerable functional redundancy that characterises E3 ligases (69).

When interpreting our bar-seq data, we reasoned that since null mutants were more likely to exhibit decreases in proportional representation as the experiments progressed, our analysis should be limited to the identification of loss-of-fitness phenotypes. This is because decreases in the relative abundance of a subset of null mutant lines in the population would lead to an increase in the proportional representation of the remaining null mutant lines, potentially mimicking gain-of-fitness phenotypes. Despite this limitation, the bar-seq approach allowed us to identify numerous fitness phenotypes associated with the promastigote and amastigote life cycle stages. In particular, loss-of-fitness was identified for more than one interval between promastigote time points for Δ*ubc9*, hinting at a role for SUMOylation (UBC9), and *Δhect2* and *Δhect12*, for HECT E3-mediated ubiquitination, in promastigote growth and/or survival. These genes were not absolutely required for survival, however, given our ability to detect Δ*ubc9*, Δ*hect2* and Δ*hect12* in the metacyclic promastigote samples. Also of interest was the high degree of correlation between the data from the axenic amastigote, macrophage and mouse infection experiments, supporting previous findings that only small transcriptomic differences exist between axenic and intracellular amastigotes (3). The strong phenotypes observed for Δ*ubc1/cdc34*, Δ*ubc2*, Δ*uev1* and Δ*hect2* in the amastigote stages suggest an important role for these genes in the successful transformation from promastigote to amastigote. Since Δ*hect2* also showed loss-of-fitness in the promastigote stage, the effect of HECT2 deletion on cell survival/proliferation may be a more general one. For example, if the function of human UBE3C is shared with HECT2, then the build-up of harmful, incompletely-degraded proteasome substrates during the differentiation process could explain the requirement for HECT2 in amastigotes and the more subtle effect of HECT2 deletion on promastigotes (32). Notably, the observed requirement for UBC1/CDC34 during *L. mexicana* mouse infection mirrors the finding that TbCDC34 is required for infection of mice with bloodstream form *T. brucei* (25). Additionally, the human orthologues of both UBC2 and UEV1 (UBE2N and UBE2V1 respectively) have been implicated in the differentiation of various human cell types, perhaps pointing to a general role for these protein families in differentiation processes (27, 70-72). Since both ubiquitination and deubiquitination enzymes have been found to be essential in the promastigote to amastigote transition, interplay between the activities of E2/E3s (UBC1/CDC34, UBC2, UEV1 and HECT2) and DUBs (DUB4, DUB7 and DUB13) could be crucial for maintaining an optimal abundance and/or state of modification of protein targets that are required for differentiation. Δ*uba1b*, Δ*ubc14*, Δ*hect7* and Δ*hect11* were lost at later stages and Δ*ubc9* showed cumulative loss throughout the experiments, suggesting that the deleted genes are required for normal amastigote proliferation, including during mouse infection.

Our finding that UBC2 and UEV1 are highly conserved at both the sequence and structural level suggests that their function may be shared between distantly related species. For example, that UEV1 is more similar in sequence to HsUBE2V2 and ScMMS2 than to HsUBE2V1 hints at a possible function in DNA damage repair (36). Our observation that UBC2 and UEV1 form a heterodimer was not unexpected and is consistent with the similar phenotypes observed for Δ*ubc2* and Δ*uev1* in promastigote to amastigote differentiation. We also found that UBC2 is a monomer *in vitro*. This property is similar to HsUBE2N but contrasts with ScUbc13, which is a homodimer *in vitro* (41). Given that dimerisation does not affect ubiquitin transfer by another human E2, UBE2W, this difference may not be physiologically relevant (73). Our X-ray crystal structure of the UBC2-UEV1 heterodimer revealed high conservation of the UBC2 active site and UBC2-UEV1 interface, highlighting the importance of these features in the function of this complex across diverse eukaryotes.

Subsequent biochemical characterisation of UBC2 and UEV1 revealed that UBA1a and UBC2 are a functional ubiquitin E1-E2 pair. This demonstrates a level of specificity for E1-E2 interactions in *L. mexicana*, despite both UBA1a and UBA1b being more closely related to HsUBA1 than HsUBA6. That UBA1a and UBA1b appear to have unique functions is interesting since there is no obvious requirement for two ubiquitin E1s in other single-celled eukaryotes. *S. cerevisiae*, for example, has only one ubiquitin E1. A differential requirement for UBA1a and UBA1b is further supported by the likelihood that UBA1a, but not UBA1b, is an essential gene in promastigotes.

The UBC2-UEV1 heterodimer, like its *S. cerevisiae* and human orthologues, is able to form K63-linked ubiquitin chains *in vitro* (28, 36, 41, 42, 45). Most of the chains observed in our *in vitro* assays were di-ubiquitin, supporting our suggestion that UEV1 is more similar to human UBE2V2, which forms di-ubiquitin *in vitro*, than UEV-1A (isoform 2 of UBE2V1), which forms polyubiquitin chains (36). Furthermore, our structural modelling of UBC2-UEV1 in complex with donor and acceptor ubiquitins, together with the conservation of key UEV1 residues that are required for acceptor ubiquitin interaction, are consistent with a role for UEV1 in dictating K63-linked chain specificity by correctly orienting the acceptor ubiquitin (47). In contrast, the donor ubiquitin is thought to exhibit flexible positioning around the covalent Cys85 linkage (47, 74). The physiological role of ubiquitin chains generated by *Leishmania* UBC2-UEV1 is currently unknown and an important area for further investigation. However, previous research has shown that free ubiquitin chains are present in both *S. cerevisiae* and human cells and that their levels (in *S. cerevisiae*) increase following heat shock, DNA damage or oxidative stress (75). Furthermore, unanchored K63-linked chains generated by the human E3 ligases TRAF6 and TRIM32 have been shown to interact with and activate protein kinases (76, 77) and unanchored K48-linked chains can inhibit the proteasome (78), demonstrating that free ubiquitin chains can perform regulatory functions. Curiously, *S. cerevisiae* HUL5, the E3 ligase partly responsible for free chain formation upon stress induction (75), is related to *L. mexicana* HECT2 (45% query cover, 32.6% identity, E value: 5e^-47^). This raises the possibility that HECT2 is involved in the response to environmental stresses that trigger the promastigote to amastigote transition and may explain the requirement for HECT2 in amastigotes. Exploring the potential interaction between UBC2-UEV1 or HECT2 and protein kinases involved in stress responses is an interesting avenue for further study. Additionally, the identification of UBC2 and UEV1 in an interactome of *L. mexicana* DUB2, which is able to cleave K63-linked diubiquitin *in vitro*, suggests a possible interplay between these proteins in the regulation of K63-linked ubiquitin chains (8).

For our *in vitro* experiments, human BIRC2 and RNF8 provided useful tools for examining the activity of UBC2 in the absence of available *Leishmania* E3s. Both BIRC2 and RNF8 were able to form polyubiquitin chains (non-K63-linked) in a UBC2-dependent manner. Intriguingly, UEV1 effectively inhibited this reaction by switching UBC2 activity towards unanchored K63-linked di-ubiquitin formation. This ability of UEV1 could allow UBC2 to flip between direct (covalent attachment of ubiquitin to substrates) and indirect (unanchored ubiquitin chain binding to regulated proteins) mechanisms of regulating other proteins as well as between sets of protein targets, depending on the availability of UEV1. The proposed role for UEV1 in specifying K63-linked chain formation is further supported by the observation that polyubiquitin chains produced in the presence of UBC2 and BIRC2 or RNF8 are not K63-linked. In support of our finding that UBC2 acts alone with RNF8, human UBE2V1 and UBE2V2 have been shown to be dispensable for the function of UBE2N and RNF8 in DNA damage signalling (79). Conversely, the formation of polyubiquitin chains on human CHIP was seen in the presence of both UBC2 and UEV1. In this example, K63-linked ubiquitin chains could be formed in a RING-dependent manner similar to that described for rat RNF4, where the RING domain stabilises the positions of human UBE2N holds the donor ubiquitin in a ‘folded-back’ conformation poised for nucleophilic attack by K63 of the acceptor ubiquitin bound to UBE2V2 (74). The ability of UBC2-UEV1 to extend ubiquitin chains on CHIP illustrates the potential for UBC2-UEV1 to act in coordination with other E2s in *L. mexicana*, although an E2 with such a role has yet to be identified.

Previous studies have investigated the importance of DUBs and the parasite proteasome at various stages of the *Leishmania* life cycle (8, 15). Supplementing this, our study explores the requirement for selected E1, E2 and E3 enzymes across the life cycle of *Leishmania*. Our detailed investigation of UBC2 and UEV1, identified as essential in amastigotes, demonstrates a high level of conservation at both the structural and functional level. Consequently, our finding that the nature of UBC2 activity can be regulated by UEV1 has implications for orthologous proteins in other species.

## Materials and Methods

### Bioinformatic identification of ubiquitination genes

Ubiquitin E1-activating enzyme, E2-conjugating enzyme and E3 ligase genes were identified in the *L. mexicana* genome by performing Interpro and PFAM domain searches in TriTrypDB (https://tritrypdb.org/tritrypdb/). The following Interpro and PFAM identification codes were used: IPR018075, PF10585, IPR019572, IPR028077, IPR000608, IPR000569, PF00632, IPR002867, PF01485, IPR001841, IPR011016, IPR003613 and PF04564. UniProt (https://www.uniprot.org/) was also used to search for genes annotated with the terms “HECT” or “RBR”. Protein BLAST searches were used to determine the percentage query cover, percentage identity and E value for two gene sequences.

### *Leishmania* culture

*L. mexicana* (MNYC/BZ/62/M379) promastigotes were grown in HOMEM (Gibco) supplemented with 10% v/v heat-inactivated Fetal Bovine Serum (FBS) (Gibco) and 1% v/v Penicillin/Streptomycin (Sigma-Aldrich) at 25°C. Typically, cells were split around twice a week. Selection drugs were added to the medium as appropriate: 10 µg mL^-1^ blasticidin (InvivoGen), 40 µg mL^-1^ puromycin (InvivoGen), 10-15 µg mL^-1^ G418 (InvivoGen), 50 µg mL^-1^ hygromycin (InvivoGen) and 50 µg mL^-1^ nourseothricin (Jena Bioscience).

### Null mutant library generation

Null mutants were generated using the CRISPR-Cas9-based approach as previously described (8, 31). Primer sequences to allow amplification of the single guide DNAs (sgDNAs) and repair cassettes for gene deletion were designed using a web tool (http://www.leishgedit.net/Home.html). Additionally, a T7 polymerase promoter sequence, 10 nt linker and 12 nt unique barcode were inserted into the 5’ end of the upstream forward primer in the following order: 5’-TAATACGACTCACTATAAAACTGGAAGXXXXXXXXXXXX-3’, where X represents the barcoded region.

PCR reactions for cassette amplification contained 30 ng of plasmid (pTBlast_v1, pTPuro_v1 or pTNeo_v1, sequences available from leishgedit.net), 0.2 mM dNTPs, 2 µM each of forward and reverse primer, 1U Q5 DNA Polymerase (NEB), 1x Q5 reaction buffer (NEB) and distilled water to make the volume up to 40 µL. The reverse primer used for generating all guides was aaaagcaccgactcggtgccactttttcaagttgataacggactagccttattttaacttgctatttctagctctaaaac. The PCR was run with the following settings: 94°C for 5 min, 45 cycles of 94°C for 30 s, 65°C for 30 s and 72°C for 2 min 15 s and 72°C for 7 min. For the sgRNAs, PCR reactions were set up in a similar manner but with a total volume of 20 µL. The PCR program used was 98°C for 30 s, 35 cycles of 98°C for 10 s, 60°C for 30 s and 72°C for 15 s and 72°C for 10 min.

Mid-log phase *L. mexicana* Cas9 T7 procyclic promastigotes were transfected either with whole PCR reactions or 5 µL of DNA purified using the QIAquick PCR Purification Kit (Qiagen). 8 x 10^6^ log phase cells were prepared by spinning down (1,000 x g for 10 min), washing with 1 x PBS and resuspending in 1 x Cytomix (66.7 mM Na_2_HPO_4_, 23.3 mM NaH_2_PO_4_, 5 mM KCl, 50 mM HEPES and 150 µM CaCl_2_, pH 7.3) or P3 solution from the P3 Primary Cell 4D-Nucleofector X Kit (Lonza). Next, cells were pulsed twice using the Nucleofector 2B device (Lonza) and the X-001 program or using the Amaxa 4D-Nucleofector (Lonza) and the FI-115 programme (for the Cytomix and P3 solutions respectively) and then placed into 5 mL of HOMEM media with 20% FBS and 1% Penicillin/Streptomycin. As a negative control, the parental cell line was transfected with water in place of DNA.

Following recovery of the cells at 25°C, appropriate antibiotics were added (10 µg mL-1 blasticidin [InvivoGen], 40 µg mL-1 puromycin [InvivoGen], 15 µg mL-1 G418 [InvivoGen]) to select for transfectants. Selection was performed either on a population level (10 mL containing the total transfected population) or along with cloning into 96-well plates at 1:6, 1:66 and 1:726 dilutions. Dilutions were carried out in HOMEM supplemented with 20% v/v FBS and 1% v/v Penicillin/Streptomycin.

To extract genomic DNA for analysis, 500 µL-5 mL of mid-log phase promastigotes were spun down (1,000 x g for 10 min), washed once in 1 x PBS and processed using the Qiagen DNeasy Blood and Tissue Kit and the manufacturer’s protocol. For genotype analysis, PCR reactions were set up with 1 μL of genomic DNA and either Q5 DNA Polymerase (NEB) or LongAmp Taq (NEB) with the manufacturer’s protocols.

### Bar-seq screen

The bar-seq screen was performed as described previously in Damianou *et al*., 2020 (8); the raw data are available in S4 Table. Statistical analyses were performed between adjacent experimental time points using paired t-tests and the Holm-Šídák method in GraphPad Prism 8.

### Cell viability assay

*L. mexicana* cultures were grown to stationary phase (>1 x 10^7^ mL^-1^) and resuspended at 1 x 10^6^ cells per mL in amastigote medium (Schneider’s Drosophila medium [Gibco], 20% FBS [Gibco] and 15 µg mL-1 HEMIN [Sigma], adjusted to pH 5.5). 200 µL cell samples were prepared in sextuplicate in 96-well plates and included the parental cell line (Cas9 T7) as a positive control. Also included were media-only, negative control samples. At 0 h, 48 h and 120 h, 20 µL of 125 µgmL^-1^ resazurin (in 1 x PBS) was added to sample wells and the plate incubated at 37°C for 8 h. The fluorescence at 590 nm was then read using the POLARstar Omega Plate Reader (BMG Labtech). Relative viability was calculated for each sample by averaging fluorescence readings across the 6 replicates, subtracting the average for the negative control and then dividing by the value obtained for the Cas9 T7 positive control.

### Sequence alignments

*S. cerevisiae* sequences were obtained from UniProtKB (80), human sequences from UniProtKB or NCBI and *L. mexicana* and *T. brucei* sequences from TriTrypDB (81). Sequence alignments were performed using T-Coffee (82) and structural annotation using ESPript 3.0 (83).

### Protein expression and purification

*UBA1a* (LmxM.23.0550), *UBC2* (LmxM.04.0680) and *UEV1* (LmxM.13.1580) genes were codon-optimised for *E. coli* expression and synthesised by DC Biosciences. Genes were amplified by PCR and inserted into an expression vector (Protein Production Facility, University of York) in frame with an N-terminal His tag and Im9 solubility tag (37) using In-Fusion cloning (Takara). Primers used for gene amplification were 5’-TCCAGGGACCAGCAATGCTTTCTGAGGAAGAGCAAAAAC-3’ and 5’-TGAGGAGAAGGCGCGTTAAAAGCGATAGCGGTAGCGGATG-3’ for UBA1a, 5’-TCCAGGGACCAGCAATGTTGACCACTCGTATCATTAAGG-3’ and TGAGGAGAAGGCGCGTCATGGTTTGGCGTACTTACGAG-3’ for UBC2 and 5’-TCCAGGGACCAGCAATGGTCGAGGTTCCGCGC-3’ and 5’-TGAGGAGAAGGCGCGTCAGTAGGTACTACCCTCC-3’ for UEV1. Insert integration and sequence were confirmed by DNA sequencing (Eurofins). Plasmids were then transformed into BL21-Gold (DE3) cells (Agilent Technologies). Cells were grown overnight in 5 mL of LB medium with 25 µgmL^-1^ kanamycin and used to seed 500 mL cultures for growth at 37°C. When an optical density of around 0.5 at 600 nm was reached, cultures were equilibrated to 20°C and 1 mM of isopropyl 1-thio-β-D-galactopyranoside was added to induce recombinant protein production. Cultures were then grown for a further 24 h at 20°C.

The standard purification procedure was as follows: bacterial pellets from 500 mL cultures were resuspended in 25 mL buffer A (20 mM NaH_2_PO_4_·2H_2_O, 20 mM Na_2_HPO_4_·12H_2_O, 0.3 M NaCl, 30 mM imidazole and 5 mM β-mercaptoethanol, pH 7.4), DNase-treated and homogenised using a cell disruptor (Constant Systems Ltd). The lysate was centrifuged (35,000 x g, 10 min, 4°C), filtered and loaded onto a HisTrap Fast Flow Crude column (GE Healthcare). After column washes with buffer A, a gradient of buffer B (20 mM NaH_2_PO_4_·2H_2_O, 20 mM Na_2_HPO_4_·12H_2_O, 0.3 M NaCl, 0.5 M imidazole and 5 mM β-mercaptoethanol, pH 7.4) was applied to elute bound protein. His-Im9 tag cleavage was then carried out using at least one tenth HRV 3C protease (Protein Production Facility, University of York) to sample protein (mg). This was followed by imidazole removal using either overnight dialysis at 4°C in buffer C (20 mM NaH_2_PO_4_·2H_2_O, 20 mM Na_2_HPO_4_·12H_2_O, 0.3 M NaCl and 5 mM β-mercaptoethanol, pH 7.4) or a desalt column. Removal of the His-Im9 tag leaves an additional 3 amino acids (Gly, Pro, Ala) at the N-terminus. Following cleavage, the sample was reapplied to the HisTrap column and application of buffer A and B used to separate the tag from the protein of interest. Sample fractions were concentrated and size-exclusion chromatography (SEC) carried out using HiLoad 16/600 Superdex 200 pg or 75 pg columns (GE Healthcare) for UBA1a and UBC2 or UEV1 respectively. Buffer used for SEC contained 50 mM HEPES and 150 mM NaCl with either 2 mM DTT (UBC2 and UEV1) or 1 mM TCEP (UBA1a). Fractions containing purified protein were identified by SDS-PAGE analysis, pooled, concentrated and stored at -80°C.

For the UBC2 and UEV1 samples used for SEC-MALLS, purification was carried out as described above but with the following minor changes: buffers A-C did not contain β-mercaptoethanol, UBC2 was not purified by SEC and the buffer used for SEC of UEV1 was 25 mM Tris-HCl, 150 mM NaCl, pH 8. Storage was in the final purification buffers plus 1 mM DTT.

### Size-exclusion chromatography multi-angle laser light scattering (SEC-MALLS)

Prior to loading, mixtures of UBC2 and UEV1 were prepared as required and incubated on ice for 30 min. 120 µL of each sample was then loaded onto a Superdex 200 HR10/300 gel filtration column (Sigma-Aldrich) and run in 25 mM Tris-HCl, 150 mM NaCl and 1 mM DTT, pH 8. Light scattering and refractive index measurements were taken using a Dawn Heleos II and Optilab rEx detector (Wyatt) respectively. Concentrations of samples loaded were between 1.2 and 2 mg mL^-1^.

### Crystallisation and structure determination

For crystallisation experiments, UBC2 and UEV1 were mixed in a 1:1 molar ratio to a final concentration of 6.6 mg mL^-1^ in buffer containing 50 mM HEPES, 150 mM NaCl and 2 mM DTT and incubated on ice for 30 min. Crystallisation conditions were screened (PACT premier HT-96 screen, Molecular Diagnostics) in a 96-well sitting drop format. Drops consisting of 150 nL of protein and 150 nL of reservoir solution were mixed and incubated above 100 µL of reservoir solution. Crystals appeared after two days at 25°C in drops prepared with a reservoir solution consisting of 0.1 M Bis-Tris propane, pH 7.5, 0.2 M sodium formate and 20% PEG. A single crystal was captured in a fine nylon loop, cryo-cooled in liquid nitrogen, and sent for data collection at the Diamond Light Source (Beamline I03). The diffraction data, extending to a nominal resolution of 1.7 Å, were processed using the 3dii pipeline in *xia2* (84). The crystals belonged to space group *P*2_1_ with two UBC2-UEV1 heterodimers in the asymmetric unit.

The structure was solved by molecular replacement in the program MOLREP (85) implemented in the CCP4i2 interface (86). The search model used was the coordinate set for the human UBE2N-UBE2V2 complex (PDB ID: 1J7D). Model rebuilding and refinement were carried out using iterations of the programs Buccaneer (87) Refmac5 (88, 89) and Coot (90) in CCP4i2 (86). The electron density maps were of good quality allowing the confident tracing of the protein chains in the two heterodimers (AB and CD) with the exception of residues in the α1-β1 loop of UEV1 chains (B and D) where the maps were of poorer quality such that residues Gly22 and Ser23 could not be built in Chain D. It is assumed that this region of the structure has higher mobility. Data collection and refinement statistics are given in S3 Table. The coordinates and structure factors for the *L. mexicana* UBC2-UEV1 complex are available in the Protein Data Bank (PDB ID:6ZM3).

### Structure analysis

Superposition of structures and RMSD determination were performed using “superpose structures” in CCP4mg (91). UBC2-UEV1 interface analysis was performed using PISA (version 1.52) (92).

### Thioester formation assay

Reactions contained 300 nM UBA1a, 2.5 µM E2, 20 µM human ubiquitin (Boston Biochem) and 5 mM ATP as indicated in 40 µL ubiquitination assay buffer (50 mM HEPES, pH 7.5, 100 mM NaCl, 10 mM MgCl_2_ and 2 mM DTT). Reactions were incubated for 0-10 min at room temperature and then quenched with sample buffer with or without reducing agent. Samples containing reducing agent were heated at 90°C for 5 min. Samples were run on an SDS-PAGE gel and stained with InstantBlue Coomassie Protein Stain (Expedeon). For the UBE2W thioester assay, bovine ubiquitin (Ubiquigent) was used in place of human ubiquitin.

### Di-ubiquitin formation assay

Reactions contained 100 nM UBA1a, 2.5 µM of UBC2 and UEV1, 100 µM of human wild-type (Boston Biochem) or K63R ubiquitin (2B Scientific) and 5 mM ATP as indicated in 40 µL of ubiquitination assay buffer. Samples were incubated at 37°C for between 0 and 90 min as indicated. For the inhibitor assay, 0-50 µM of NSC697923 (Abcam) was pre-incubated with UBC2 and UEV1 in reaction buffer for 15 min at room temperature prior to addition of UBA1a and ATP. The reaction was then incubated at 37°C for 90 min. The final DMSO concentration in these reactions was 0.5%. Following incubation, reducing sample buffer was added and the samples heated at 90°C for 5 min. Samples were separated by SDS-PAGE and Western blotting carried out using a mouse mono- and polyubiquitinated ubiquitin conjugate (Ubiquigent) or mouse Ub-K63 (ThermoFisher) antibody with HRP-conjugated anti-mouse secondary antibody (GE Healthcare or Promega). In addition to the experimental samples, 100 ng of K63 di-ubiquitin positive control (Ubiquigent) was loaded where indicated.

### E3 cooperation assay

Reactions were prepared with 100 nM UBA1a, 2.5 µM E2, 1 µM human E3, 100 µM ubiquitin (Boston Biochem) and 5 mM ATP as indicated in ubiquitination assay buffer in 40 µL total reaction volume. BIRC2, RNF8 and (N-terminally truncated) HUWE1 were all GST-tagged and sourced from Ubiquigent. Samples were then incubated at 30°C for 1 h prior to SDS-PAGE and Western blotting with a mouse mono- and polyubiquitinated ubiquitin conjugate (Ubiquigent), mouse Ub-K63 (ThermoFisher) or rabbit anti-GST (Abcam) antibody with HRP-conjugated anti-mouse (GE Healthcare) or HRP-conjugated anti-rabbit (GE Healthcare) secondary antibody as appropriate.

### CHIP priming and extension assay

Reactions were prepared with 0.1 µM UBA1a, 2.5 µM 6His-tagged UBE2W (Ubiquigent), 1 µM CHIP (Ubiquigent), 0.1 mM human ubiquitin (Boston Biochem) and 2 mM ATP (Sigma-Aldrich) in ubiquitination assay buffer in 50 µL total reaction volume. Samples were incubated at 30°C for 1 h prior to SDS-PAGE and Western blotting with a mouse mono- and polyubiquitinated conjugate (Ubiquigent) or rabbit CHIP antibody (Calbiochem) with HRP-conjugated anti-mouse (GE Healthcare) or HRP-conjugated anti-rabbit (GE Healthcare) secondary antibody as appropriate.

## Supplementary methods

### Thioester formation assay

Reactions contained 6 pmol E1, 2.5 µM E2, 2 nmol human ubiquitin (Boston Biochem) and 2 mM ATP as indicated in 20 µL ubiquitination assay buffer. Reactions were incubated for 30 min at 30°C. Sample buffer with or without reducing agent was then added and the samples containing reducing agent heated at 90°C for 5 min. Samples were run on an SDS-PAGE gel and stained with InstantBlue Coomassie Protein Stain (Expedeon). Recombinant *L. mexicana* UBA1b was donated by Daniel Harris, University of Glasgow.

## Acknowledgements

We would like to thank Ubiquigent for providing a 3-month internship for RB where the biochemical characterisation of UBC2-UEV1 was carried out. Specifically, we would like to thank Jane Hilton and Sheelagh Frame for technical support and Jason Brown for reviewing the manuscript. Additionally, we thank Daniel Harris (University of Glasgow) for donating the recombinant *L. mexicana* UBA1b and Farid El Oualid (UbiQ) for valuable discussions. We also thank Diamond Light Source for beamline access to I03 (Proposal No MX-18598-30) that contributed to the results presented here, Marek Brzozowski for crystal-harvesting and valuable discussions, Johan Turkenburg for help with data processing and Eleanor Dodson for advice on structure solution and refinement.

## Figure legends

**S1 Table**. *Summary of L. mexicana ubiquitination genes*.

**S2 Table**. *Individual viability assay of Δubc1, Δubc2, Δuev1 and Δhect2 during axenic amastigote differentiation*. Raw fluorescence (590 nm) readings and relative viabilities compared to the Cas9 T7 cell line are included.

**S3 Table**. *Data collection and refinement statistics for UBC2-UEV1 crystal structure*.

**Supplementary figure 1**. *Alignments of L. mexicana and H. sapiens protein sequences*. **A** ubiquitin E1-activating enzyme, **B** ubiquitin E2-conjugating enzyme and **C** E3 ligase HECT domain alignments. Sequences obtained from TriTryDB (*L. mexicana*) and NCBI or UniProt (*H. sapiens*) were aligned using T-Coffee. Red boxes indicate amino acid identity, red characters show similarity within the highlighted group and blue frames highlight similarity across groups. Black bar indicates the position of the conserved HPN motif in E2 genes and the black triangles highlight the conserved catalytic cysteine residues in all classes of protein. *L. mexicana* genes are indicated with the prefix Lm and *H. sapiens* genes with the prefix Hs. Where no prefix is given, sequences belong to *L. mexicana*.

**Supplementary figure 2**. *Confirmation of the null mutant library*. **A** Schematic of PCRs performed to identify null mutants from heterozygotes or mutants with additional gene copies. Primers were designed to amplify regions within the gene of interest (red primers) and between the 5’-UTR of the gene and the blasticidin resistance cassette of the edited DNA (black primers). **B** PCRs performed on genomic DNA from parental (P) and mutant (M) cells are shown for genes that were deleted successfully. Due to the large size of *HECT3* (18.6 kbp), only the first 8 kbp of the gene was targeted for editing. The *HECT3* gene PCR was designed to amplify within the targeted region. **C** PCRs performed on genomic DNA from parental (P) and mutant (M) cells are shown for genes that could not be deleted (no mutant clones were obtained for UBC13). **D** Relative viability of Δ*ubc1*, Δ*ubc2*, Δ*uev1* and Δ*hect2* compared to the parental Cas9 T7 line during promastigote to amastigote differentiation. Time elapsed since the initiation of differentiation is marked on the x-axis. Data are an average of two biological replicates, each with 6 technical replicates.

**Supplementary figure 3**. *Structural analysis of the UBC2-UEV1 heterodimer*. **A** Superposition of chains A and B for UBC2-UEV1 (dark green and dark blue, PDB ID: 6ZM3), HsUBE2N-UBE2V2 (darker green and darker blue, PDB ID: 1J7D) and ScUbc13-Mms2 (light green and light blue, PDB ID: 1JAT). Locations of the N- and C-termini are shown. **B** Zoom-in of the interface between UBC2 and UEV1 showing residues thought to contribute most significantly to complex formation according to analysis in the program PISA (92). Residues are labelled in black for UBC2 and blue for UEV1. Residues are coloured by atom (red, oxygen; blue, nitrogen). Hydrogen bonds are denoted by dashed lines. **C** Superposition of UBC2-UEV1 onto the structure of the UBE2N-UBE2V2-Ub complex (PDB ID: 2GMI) (47) showing UBC2 (green) and UEV1 (blue) as space fill models and the positions of acceptor and donor ubiquitins (orange ribbons) obtained from the UBE2N-UBE2V2-Ub structure. C85 is represented by black (carbon) and yellow (sulfur) spheres. K63 of ubiquitin is highlighted as cylinders coloured by atom (blue, nitrogen).

**Supplementary figure 4**. *UBA1b cannot transfer ubiquitin to UBC2 in vitro*. **A** Alignment of *L. mexicana* and *H. sapiens* ubiquitin protein sequences. Red boxes indicate amino acid identity, red characters show similarity within the highlighted group and blue frames highlight similarity across groups. **B** UBA1b and ubiquitin were incubated with UBC2, UEV1 and ATP as indicated in ubiquitination assay buffer for 30 min at 30°C. Samples were treated with either reducing or non-reducing sample buffer and visualised by SDS-PAGE with InstantBlue stain.

